# Architecture of intact mammalian pyruvate dehydrogenase complex revealed by *in-situ* structural analysis

**DOI:** 10.1101/2024.02.01.578502

**Authors:** Chen Wang, Cheng Ma, Shenghai Chang, Hangjun Wu, Chunlan Yan, Jinghua Chen, Yongping Wu, Shaoya An, Jiaqi Xu, Qin Han, Yujie Jiang, Zhinong Jiang, Haichun Gao, Xing Zhang, Yunjie Chang

## Abstract

The multi-enzyme pyruvate dehydrogenase complex (PDC) links glycolysis to the citric acid cycle by converting pyruvate into acetyl-coenzyme A and is essential for cellular energy metabolism. Although individual components have been characterized, the structure of intact PDC remains unclear, hampering our understanding of its composition and multi-step catalytic reactions. Here, we report the *in-situ* architecture of intact PDC within mammalian mitochondria by cryo-electron tomography. The organization of peripheral E1 and E3 multimers varies substantially among PDCs. We discovered 1-46 E1 heterotetramers surrounding each PDC core, and up to 12 E3 dimers locating primarily along the pentagonal openings. Furthermore, we observed dynamic interactions of the substrate translocating lipoyl domains (LDs) with both E1 and E2, implicating the mechanism of substrate channeling. Our findings reveal the intrinsic dynamics of PDC peripheral compositions, suggesting an additional model in which the number of assembled E1 heterotetramer, E3 dimer, and functional LDs of PDC may be regulated to further control its catalytic activity.

---

The pyruvate dehydrogenase complex (PDC) is a multi-subunit complex that is essential for oxidative pathway and cellular energy metabolism. It links glycolysis to the citric acid cycle, and the biosynthesis of fatty acids and steroids by catalyzing the oxidative decarboxylation of pyruvate into acetyl-coenzyme A (acetyl-CoA) ^1–4^. Due to its critical role in metabolism, PDC is associated with many major diseases, such as neurodegenerative disorders, metabolic acidosis, diabetes, and cancer^5–8^.

The PDC, along with the 2-ketoglutarate dehydrogenase complex (OGDC) and the branched-chain α-keto acid dehydrogenase complex (BCKDC), belongs to the α-keto acid/2-oxo-acid dehydrogenase (OADH) family, the largest cellular enzyme system with the molecular weight ranging from 4 to 10 million Da (MDa) ^9^. The PDC is composed of multiple copies of three catalytic enzymes: pyruvate dehydrogenase (E1; EC1.2.4.1), dihydrolipoyl transacetylase (E2; EC 2.3.1.12) and dihydrolipoyl dehydrogenase (E3; EC 1.8.1.4) ^10^. E1 is a **α**2 dimeric in Gram-negative bacteria and a **α**2**β**2 heterotetrameric in Gram-positive bacteria and eukaryotes^11–13^. E2 is a multidomain protein consisting of 1-3 lipoyl domains (LDs), a peripheral subunit-binding domain (PSBD) and an inner catalytic (IC) domain (Fig. 1a). E3 is a homodimer^14,15^ and is shared among different members of the OADH family. E2 enzymes fold into a cube (24-mer, for gram negative bacteria) or pentagonal icosahedron (60-mer, for gram-positive bacteria and eukaryotes) from trimers^16–19^, serving as a scaffold for the assembly of peripheral E1 and E3 enzymes. The catalytic reaction of PDC consists of three steps (Fig. 1b). First, E1 undergoes an oxidative decarboxylation reaction and transfer of an acyl group to lipoate in mobile LD of an E2 enzyme. Then the activated E2-linked LD transports the lipoyl moiety to CoA, forming acetyl-CoA. Finally, the LD is reoxidized for another catalytic cycle by an E3 enzyme. The E3 enzyme facilitates the transfer of protons to NAD+, resulting in the formation of NADH^3^. In the catalytic reaction cycle, the LD domains function as flexible "swinging arms" to transport intermediate substrates among the active sites of the enzymes within PDC^20,21^ (Fig. 1f).

**Fig. 1.**
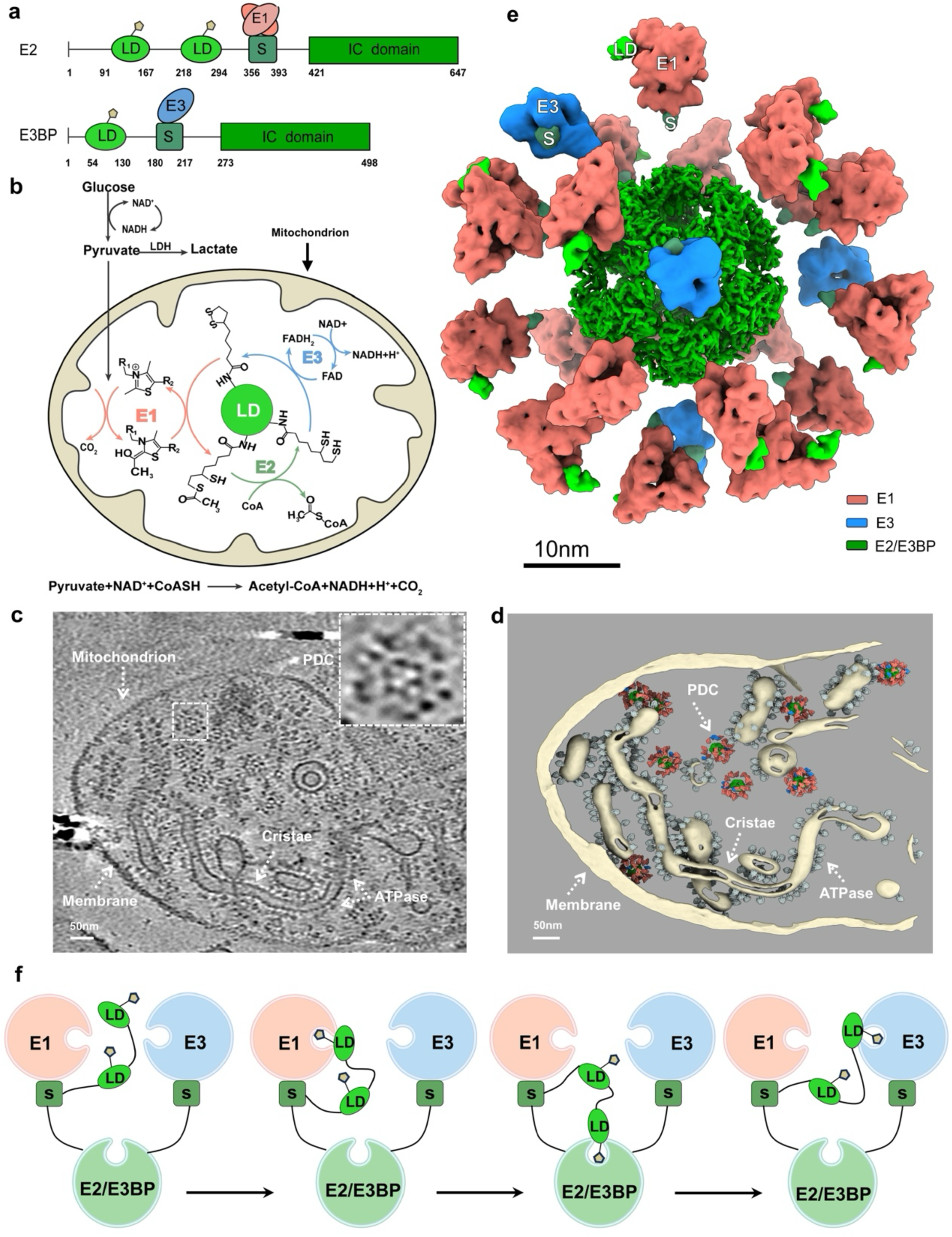
The molecular architecture and multi-step catalytic reactions of mammalian PDC. **a**, Organization of E2 and E3BP domains of mammalian PDC. E2 or E3BP is composed of one or two lipoyl domains (LDs) followed by a peripheral subunit-binding domain (PSBD) and an inner catalytic (IC) domain that are connected by flexible linkers. “S” represents PSBD. E1 or E3 binds on the PSBD of E2 or E3BP, respectively. **b,** Scheme of the multi-step reaction catalyzed by the individual components of PDC. **c,** A representative cryo-electron tomogram slice showing intact PDC particles in a mitochondrion. The insertion shows the zoomed in view of one PDC particle. **d,** Three-dimensional surface view of the tomogram shown in **c**. **e,** Average structure of mammalian PDC, with 21 E1 and 4 E3 surrounding the 60-mer E2 core (green: E2, light coral: E1, blue: E3, lime: LD, teal: S). See also Supplementary Video 1. **f,** A putative scheme (based on Arjunan et al^21^) of the substrate channeling mechanism in the catalytic cycle of PDC. Only one copy of each enzyme is shown for simplicity. From left to right: the pre-catalytic state of PDC, the first, second and third reaction step of the PDC catalytic cycle, respectively. During the catalytic cycle, the LD interacts with each enzyme sequentially to transport the substrate.

Previous studies have characterized the structures of E1, E2 and E3, and their stoichiometry, catalytic processes, and regulatory mechanisms of the PDC^11–19,22–33^. However, the majority of solved structures to date are individual components of the PDC, thus preventing the full understanding of how individual enzymes of PDC collaborate to perform catalyzes as a whole. Recently, several structures of endogenous PDC from bacteria, fungi, and bovine, with special focus on the PDC core, were obtained by cryo-electron microscopy single-particle analysis (cryo-EM SPA) ^31,34–38^ and provide valuable insights into the interactions between different components of the PDC. The interaction interface of LD and E2 IC domain was revealed from the resting state E2, which was obtained by growing *Escherichia coli* in a minimal medium supplemented with succinate instead of glucose^36^. Compared to the prokaryotic counterpart, the eukaryotic PDC has additional dihydrolipoamide dehydrogenase-binding protein (E3BP) ^39^, which is structurally very similar to E2 (Fig. 1a) and localizes at the PDC core. In fungi, E3BP was found to reside within the PDC core as four separate complexes^34,35,38^. While for mammalian PDC, multiple copies of E3BP substitute part of E2 and form an icosahedral E2/E3BP core^23^, with the stoichiometry of E2 and E3BP remains elusive. Nonetheless, these structures exhibited limited resolutions for peripheral densities and were unable to distinguish E1 or E3, hampering our understanding of the overall PDC structure. Moreover, how the substrate intermediates are channeled by LDs to other PDC components during catalysis also remains elusive.

Characterizing the OADH family complex in its intact state through crystallization or purification remains challenging due to the complexity of its assembly and the highly mobile E2-linked lipoyl "swinging arms", which play a central role in the multi-step catalytic mechanism. In addition, previous studies have suggested that PDCs display intrinsic heterogeneity, lacking uniform structures with defined stoichiometry and subunit organization^31,34,40,41^. Unraveling the molecular architecture of intact PDCs is indispensable for a comprehensive understanding of its catalytic mechanisms. Here, we determined the *in-situ* structure of intact PDC from the mitochondria of porcine hearts by cryo-electron tomography (cryo-ET). Our results show structural evidence of highly heterogeneous composition and spatial distribution of E1 and E3 multimers, providing deeper insights into collaborative mechanisms among multiple components within PDC. Furthermore, we observed new structural snapshots of the LD transfer pathway, illustrating how the swinging arms deliver the substrate-carrying prosthetic group into the active sites of E1 and E2. Overall, our work provides profound insights for understanding the regulation, assembly and collaborative catalytic reactions of multi-enzyme OADH complexes.

## *In-situ* structure of intact PDC

To characterize PDC in its native environment, we reconstructed the *in-situ* structure of PDC within mitochondria of porcine heart using cryo-ET. The mitochondrial membrane, cristae and ATPase are clearly visible in the cryo-electron tomograms (Fig. 1c-d). The cristae exhibit a conventional lamella shape, indicating the investigated regions retain the microenvironment of mitochondria. PDC particles were readily seen in the reconstructed tomograms (Fig. 1c). To get an idea about the abundance of PDC in mitochondria, we measured the volume size of the examined mitochondria in our tomograms and calculated the density of the PDC in native mitochondrial lumen. Based on our calculation, there is only one PDC particle per 2.7×10^-3^μm^3^. Considering that the volume of mitochondrion ranges from 0.04-0.08 μm^342^, the average number of PDC is approximately 15-30 per mitochondrion in heart cells.

The PDC core is found to be surrounded by a shell of protein densities, which lacks apparent symmetry and is speculated to be E1 or E3 multimers (Fig. 1c insertion). To solve the *in-situ* structure of the PDC at high resolution, we picked PDC particles from hundreds of tomograms and carried out sub-tomogram averaging analysis (Extended Data Fig. 1 and Extended Data Table 2). In the average structure, E2 folds into an icosahedral core (dark green in Fig. 1e), which agrees well with the reported mammalian PDC core structure^18,31^. The surrounding shell densities were classified, refined, and then distinguished as E1 heterotetramers or E3 dimers by comparing with the reported crystal structures (Extended Data Fig. 1). Moreover, local refinement and classification help us identified the LD and PSBD binding on the E1 or E3 multimers (Fig. 1e, details will be shown in following sections). The numbers and locations of E1 heterotetramer and E3 dimer vary across the PDC particles, which will be presented in details in the next section. On average, each PDC has twenty-one E1 heterotetramers and four E3 dimers surrounding the 60-mer E2 core (Fig. 1e and Supplementary Video 1).

## The heterogeneous organization of E1 and E3 illustrates the intrinsic flexibilities of PDC composition and assembly

E1 and E3 multimers were distinguished by 3D refinement and classification, making it possible to analyze their spatial organization. The number of E1 heterotetramers in the periphery of each PDC in mitochondria (hereafter called mitochondrial PDC) is not constant, but varies from 1 to 46, with an average of 21 E1 heterotetramers per PDC (Fig. 2a). The distribution of E1 number fits with two normal distributions (Fig. 2a). The number of E3 dimers in each mitochondrial PDC varies between 0 and 12, with an average of 4 E3 dimer per PDC (Fig. 2b). It is worth noting that each PDC has at least one E1 heterotetramer, whereas a subset of PDC particles has no E3 at all (Fig. 2b). We then further conducted statistical analysis on the quantities of E1 and E3 in the same mitochondrial PDC. The result shows a linear variation in the numbers of E1 and E3 multimers (Fig. 2c). Specifically, for each mitochondrial PDC, there was an additional assembly of E3 dimer along with five more E1 heterotetramers. In addition, the distributions of E1 and E3 multimers in the same PDC also exhibited two major proportions (Fig. 2c), with the centers locating around [7E1, 1E3] and [32E1, 6E3], which is consistent with the bimodal distribution of E1 heterotetramer (Fig. 2a). Our results show that native PDCs do not have a specific subunit stoichiometry, consistent with previous reports^35,43–45^.

**Fig. 2.**
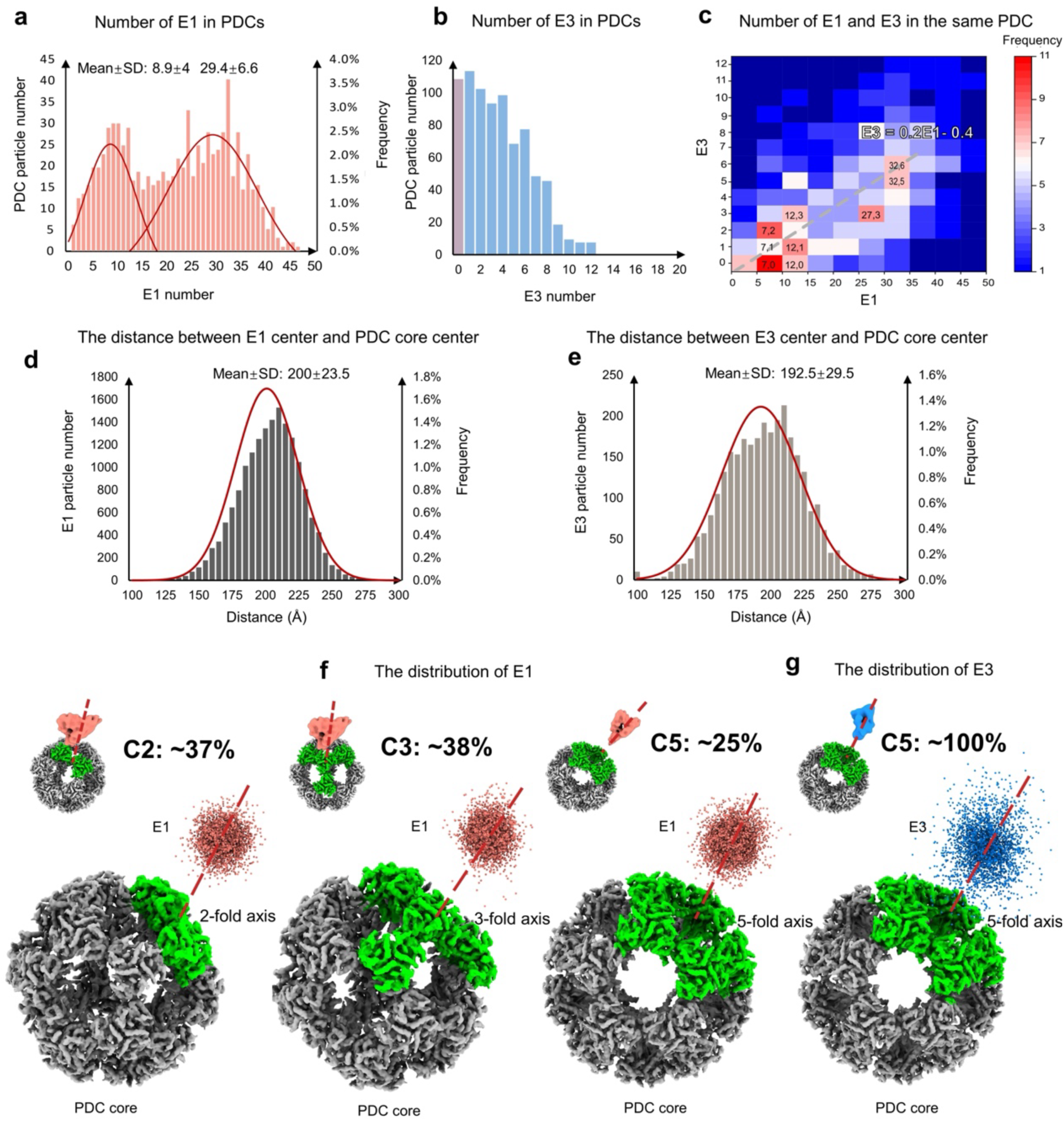
The numbers and spatial arrangements of E1 and E3 multimers in mitochondrial PDC. **a**, Distribution of the number of E1 heterotetramer in each mitochondrial PDC. SD: standard deviation. **b**, Distribution of the number of E3 dimer in each mitochondrial PDC. **c**, Distribution of the number of E1 and E3 multimers in the same PDC in mitochondria. As indicated by the grey line, the number of E3 has a linear relationship with the number of E1 (E3 = 0.2 * E1 – 0.4). **d**, Statistics of the distance between the E1 heterotetramer center and the PDC core center. **e**, Statistics of the distance between the E3 dimer center and the PDC core center. **f**, Spatial distribution of E1 heterotetramer. E1 locates along the 2-fold (left), 3-fold (middle) or 5-fold (right) axis of the icosahedral PDC core. Each light coral dot represents one E1 heterotetramer. **g**, Spatial distribution of E3 dimer. E3 primarily locates along the 5-fold axis of the PDC core. Each blue dot represents one E3 dimer. See also Supplementary Video 2.

The distances between E1/E3 multimer centers and the icosahedral PDC core were then measured to analyze the diameter of PDC. The distance between the E1 center and the PDC core center varies between 130 Å and 270 Å, with an average value and standard deviation of ∼200 Å and ∼24 Å, respectively (Fig. 2d). While the average distance between E3 center and the PDC core center is ∼193 Å, with the standard deviation ∼30 Å (Fig. 2e). Therefore, taking into account of the size of E1 and E3 multimers, the diameter of PDC typically ranges from 40 to 60 nm, with an average value of 50 nm.

The icosahedral PDC core has 2-, 3- and 5-fold symmetries. We then analyzed the spatial distributions of E1 and E3 multimers along different symmetry axes. The E1 heterotetramers scatter around the core, with minor preference locating along the 2-fold and 3-fold axes than the 5-fold axis (Fig. 2f and Extended Data Table 3). On the other hand, the E3 dimer only locates above the pentagonal opening (Fig. 2g and Supplementary Video 2).

Previous cryo-EM SPA studies using PDC purified from endogenous sources suggest that E1 and E3 multimers may form clusters surrounding the E2 core^31,35,36^, which is different from our results. To address the difference, we purified the PDC from the mitochondria of porcine hearts (hereafter called purified PDC) and frozen the samples on cryo-EM grids covered with graphene oxide. The analyses of PDC fractions using SDS-PAGE and mass spectrometry show that all components are maintained in the purified PDC (Extended Data Fig. 2 and Extended Data Table 1). Instead of doing SPA, we carried out cryo-ET investigations for the purified PDC. The tomograms show that the purified PDC encountered sample compression, resulting in fragmentation and the presence of scattered E1 and E3 subunits in the background (Extended Data Fig. 3). Some PDC particles locate around the tomogram center and possess E2 core with clear icosahedral features. However, the particles close to the graphene surfaces, looking perfect along the z-axis, are broken and have incomplete cores and periphery densities. The complete and compressed particles can be easily distinguished in 3-dimensional tomograms, but not in the projection images for SPA. We thus speculate that the mixture of the compressed particles with the complete ones increases the difficulty for the subsequent distinguishment of E1 and E3 multimers.

To further analyze the differences between the purified PDC and mitochondrial PDC, we manually picked 1209 complete particles from 256 tomograms of the purified PDC and carried out similar subtomogram averaging and statistical analysis as the mitochondrial PDC. The quantity of E1 and E3 multimers in each purified PDC varies between 1-42 and 0-11, respectively, with the average value of twenty-two E1 and three E3 multimers (Extended Data Fig. 4a and b). The distance between E1 center, E3 center and the PDC core is ∼200 Å and ∼190 Å, respectively (Extended Data Fig. 4d and e). These features are similar to those native PDCs observed within the mitochondrial lumen. However, it is notable that the number distribution of E1 heterotetramers in purified PDC fits with one normal distribution (Extended Data Fig. 4c), rather than the bimodal distribution of mitochondrial PDC. This difference could be attributed to the PDC extraction, which may lead to sample homogenization and the loss of the natural distribution observed in the mitochondrial PDC. Moreover, it further confirms that the mitochondria we observed retain their microenvironment. Together, our structural analysis here reveals the heterogeneous organization of E1 and E3 multimers, illustrating the intrinsic flexibilities of the composition and assembly of the PDC (Supplementary Videos 1 and 2).

## Binding of E1 and E3 on PSBD and the structural evidence of E1-LD interactions for alternating-active site mechanism

The PSBD subunits of the PDC core provide binding sites for E1 and E3 multimers, and the LD serves as a swinging arm to transfer substrates among different enzymes^3,20,46^ (Fig. 3a). We carried out 3D classifications to analyze these interactions of E1 and E3 with PSBD and LD. The purified PDC and mitochondrial PDC particles were combined together to increase the particle number and improve the achievable resolution (Extended Data Fig. 1). The E1 heterotetramers were classified into three main classes (Fig. 3b-d). The reported E1 crystal structure (PDB 6cfo) fit well with the main architecture of the class averages, with all E1 heterotetramers possessing an additional density close to the C-terminal region (Fig. 3b-d) and ∼82% E1 heterotetramers having additional density close to the N-terminal region (Fig. 3c-d). It was previously reported that the N terminus of E1 α subunit binds PSBD, while its C terminus interacts with LD^12,47,48^. We thus docked PSBD (PDB: 1zwv) and LD (predicted by AlphaFold2^49^, Extended Data Fig.5a) to corresponding regions, which agree well with our structures. Due to the resolution limitation, PSBD can be fitted in two different ways with opposite orientations (Fig. 3b-d and Extended Data Fig. 5b). Nevertheless, since E1 is a **α**2**β**2 heterotetramer, these two fitting strategies are thus structurally identical.

**Fig. 3.**
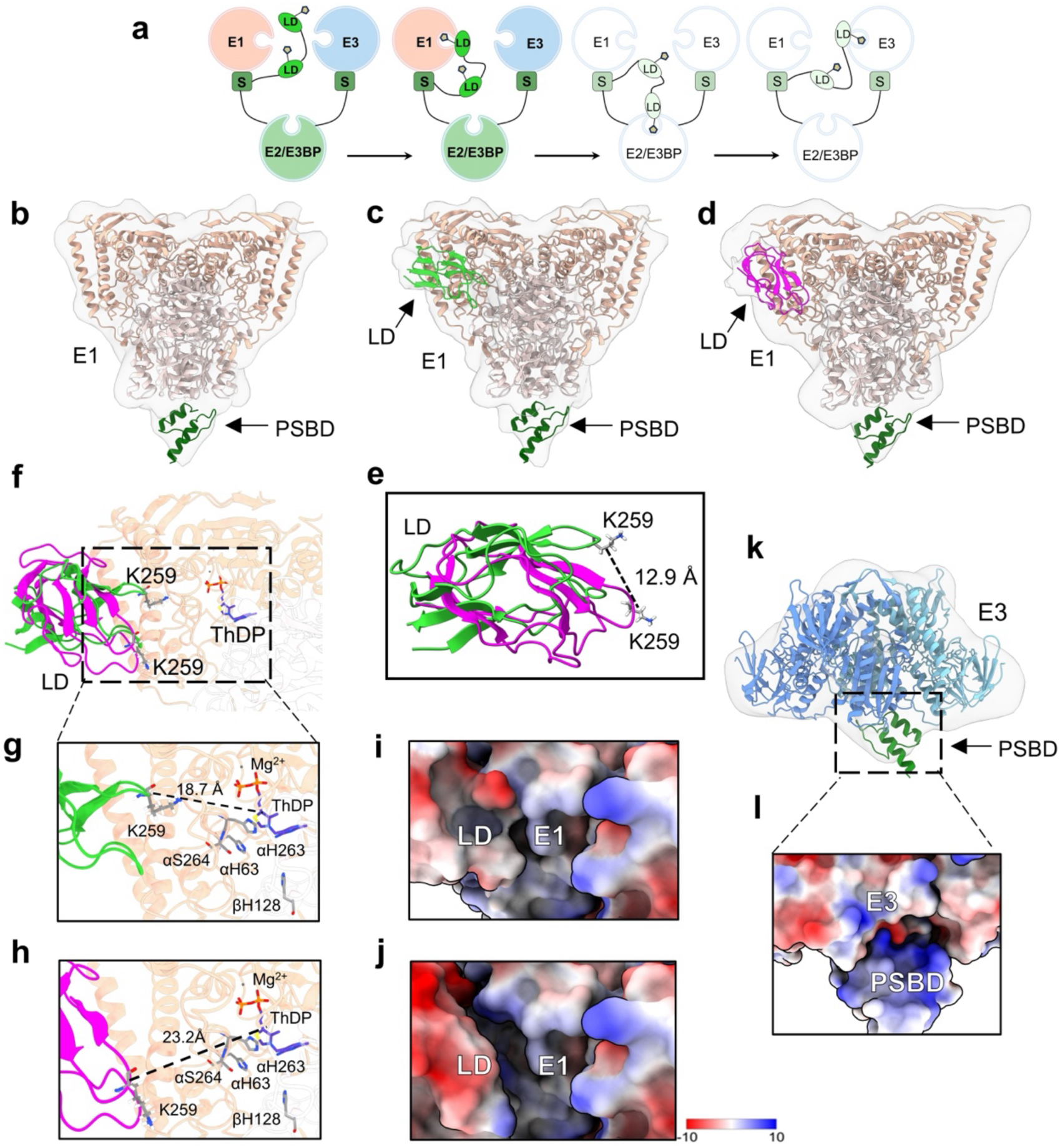
The interactions of E1 and E3 with PSBD and LD of the PDC core. **a,** The putative scheme of the substrate channeling mechanism in the catalytic cycle of PDC, as shown in Fig. 1f. The diagram highlights the pre-catalytic state and the E1-LD interactions in the first step of catalytic cycle. **b-d,** Class averages of E1 subunits. **b,** Class average showing the binding of E1 on PSBD with the rigid fitting models. PSBD locates under the E1 C-terminal region. The docking models are from the reported human source structures (PDB: 6cfo for E1 and PDB: 1zwv for PSBD). **c** and **d,** Class averages showing the interaction of E1 with PSBD and different conformations of LD. LD locates near the E1 N-terminal region. The docking model of LD was simulated from the porcine source (218-294 amino acids) using the AlphaFold2. **e,** Comparison of the two conformations of LD shown in **c** and **d**. **f,** The interaction interfaces between two conformations of LDs with E1. **g** and **h,** Enlarged view of the interaction interface shown in **f**. The distances between the lysine residue K259 of LD and the substrate ThDP is indicated. **i** and **j,** Electrostatic potential of the interaction interface between the LD and E1, corresponding to panel **g** and **h**, respectively. **k,** Average structure showing the binding of E3 on the PSBD with the rigid fitting models. Models of E3 and PSBD was obtained using the human source (PDB: 1zy8). **l,** Electrostatic potential at the interaction interface between PSBD and E3. LD: lipoyl domain, S: PSBD.

Our class averages (Fig. 3c and d) present two different binding positions of LD on E1, with its lysine residue K259 shifting ∼12.9 Å (Fig.3e). As a consequence, the lysine residue in one class average (Fig. 3g) is closer to the substrate ThDP than the other class average (Fig. 3h). Therefore, we speculate that the structure shown in Fig.3c illustrate the functional state of substrate transferring between E1 and LD, while the other structure shown in Fig. 3d represents an intermediate state of the dynamic LD binding process. In addition, the number of E1 subunits with LD binding is 4.5 times of those E1 without LD binding (Extended Data Table 3), indicating that E1 is involved in an active oxidative decarboxylation reaction. Our results also suggest that the interactions between LD and E1 primarily rely on electrostatic interactions (Fig. 3i and j). The LD presents a large negatively charged patch (E239, E254, E256, E265, E279 and D250, Extended Data Fig. 6c) that interacts with the positively charged residues of the N-terminal of the E1 α-helix (K48, K54, K307, R43, R44, R273, R275, R282, R314, H63, H92, H261and H263) (Extended Data Fig. 5c).

It is worth noting that, although the **α**2**β**2 heterotetrameric E1 has two active sites, in our results, only one active site was observed to be occupied by LD (Fig. 3c and d). Studies in biochemistry, kinetics, and spectroscopy have shown that E1 functions through an alternating active-site mechanism, also called the flip-flop mechanism, i.e., one active site is involved in the pyruvate decarboxylation process while the other catalyzes the reductive acetylation of an E2-lipoamide molecule simultaneously^50–53^. Our results offer evidence for this alternating active-site mechanism from a structural biological perspective.

Our analysis of the E3 subunits illustrates that one PSBD binds to the bridge of the E3 dimer (Fig. 3k), primarily relying on electrostatic interactions (Fig. 3l). The structures show that both the E1 heterotetramer and E3 dimer bind on one PDC-core linked PSBD. The E1 heterotetramer roughly faces the PDC core with its 2-fold axis, while the 2-fold axis of E3 dimer tilts ∼60° relative to the pentagonal opening of the PDC core (Fig. 1e and Extended Data Fig. 5d). Overall, our structure shows interactions of PSBD-E1-LD and PSBD-E3, illustrating the binding of E1 and E3 to the E2/E3BP core and the substrate transfer between E1 and LD.

## Electrostatics-based dynamic interactions between the LD and the PDC core

PDC transfers substrates between different enzymes by the LDs^16^, which are connected to the catalytic domains of the PDC core through flexible linker regions (Fig. 4a). Analysis of the interactions between LD and different enzymes of the PDC is essential for understanding the catalytic reactions. The *in-situ* structure of the PDC core was resolved by sub-tomogram averaging to analyze the interactions between LD and the PDC core. The PDC core structure was resolved to a resolution of 4.3 Å (Fig. 4b). The overall structure shows the expected 60-meric central core with icosahedral symmetry and the density matches well with the SPA resolved core structure of bovine PDC (PDB 7uom) (Fig. 4b). Noticeably, if we lower the threshold of the core structure, significant protruding densities can be found at the bridges of E2 subunits (arrow indicated in Fig. 4b), which supposed to be the LDs or flexible linkers with dynamic conformations.

**Fig. 4.**
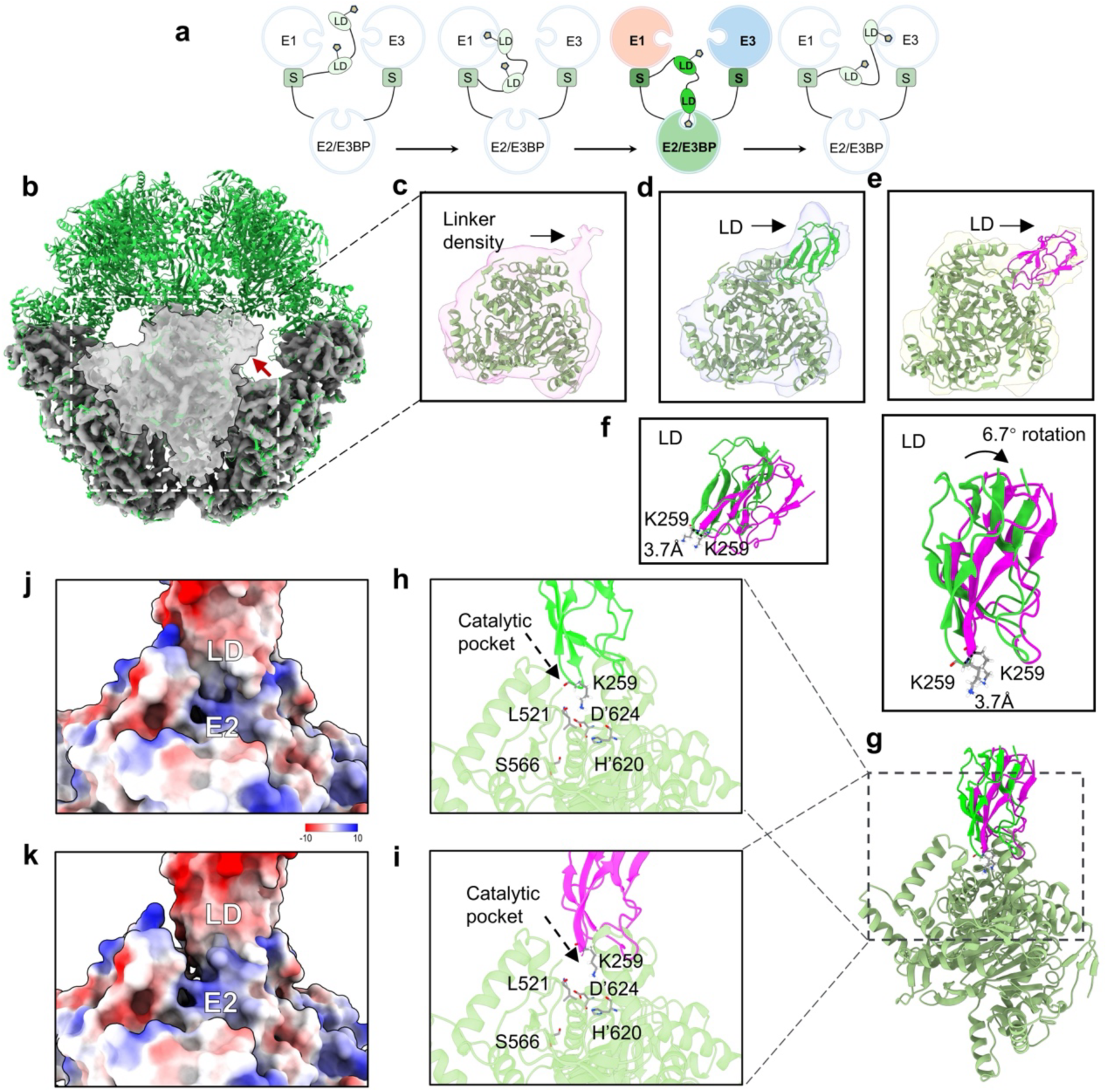
Interactions between the LD and the catalytic domain of E2. **a,** The putative scheme of the substrate channeling mechanism in the catalytic cycle of PDC, as shown in Fig. 1f. The diagram highlights the second step of the catalytic cycle, emphasizing the interactions between the LD and the PDC core in the second step of PDC catalytic cycle. **b,** 4.3 Å resolution subtomogram average structure of the inner E2/E3BP core (grey) fitting with the bovine PCD core atomic structure (green, PDB:7umo). The transparent density shows one E2/E3BP trimer with a lower threshold. The additional density indicated by the red arrow represents the extension from the PDC core. The extended densities were classified into three classes shown in **c**-**e**. **e,** One class average has a small extension density, which is supposed to be the linker between the IC domain and the LD or PSBD. E2 (PDB: 7uom) was fitted into the density map. **d** and **e,** Two class averages with large extended densities which can be fitted with E2 (PDB: 7uom) and LD. The docking model of LD was simulated from the porcine source (218-294 amino acids) using the AlphaFold2^49^. The position of LD was adjusted by rigid fitting to the density map. **f,** Comparison of the two conformations of the LD shown in **d** and **e**. **g,** The interaction interface between the LD and E2 trimer, corresponding to the structures shown in **d** and **e**. **h** and **i,** Enlarged view of the interaction interface shown in **g**. **j** and **k,** Electrostatic potential of the interaction interface between LD and E2, corresponding to **h** and **i**, respectively. LD: lipoyl domain, S: PSBD.

To better visualize the protruding densities, we performed asymmetric reconstruction of the E2 subunits. All 60 E2 subunits were extracted from each icosahedral PDC core, aligned together and then 3D classified into three classes. These class averages present significant protruding densities on the E2 trimer (Fig. 4c-e and Extended Data Fig. 6a and b). The protruding density in one class average is significantly smaller than LD and should be the extended linker (Fig. 4c). The LD (predicted by AlphaFold2^49^, Extended Data Fig.5a) fit well into the prominent densities of the remaining two classes (Figs. 4d and e). Although the locations of LD relative to E2 rotate ∼6.7° in the class averages (Figs. 4d and e), the position of the lysine residue K259 barely changes (shifts ∼3.7Å). Since the E2 active site channel is located at the interface between two catalytic domains^36^. The structures thus illustrate that the LD binds above the E2 catalytic pocket (Fig. 4g) and inserts the lipoyl moiety attached lysine residue (K259) to a funnel-shaped opening that leads to the E2 active site channel (Figs. 4h and i).

Our results suggest that the electrostatic interactions between the positively charged amino acids of LD (Extended Data Fig. 6c) and the negatively charged amino acids of E2 (R429, R430, R435, R549, K552, K547, H556 and H620, Extended Data Fig. 6d) play a crucial role in stabilizing their interactions (Fig. 4j and k). In the catalytic domain of E2, the overall surface potential around the active site channel entrance is mostly electropositive (Extended Data Fig. 6d), which could be a common electrostatic navigation mechanism across all E2 systems^36^. Moreover, the sequence alignments of the LDs in prokaryotes, fungi, and mammals show that the number of acidic amino acids in different species ranged from 12 to 15 with conserved electric properties (Extended Data Fig. 6e). This indicates a general electrostatic-based interaction between LD and E2 across species. In brief, we captured two snapshots of the previously indistinct E2-LD interactions in mammalian PDC, providing insights into the transacetylase reaction taking place within mitochondria.

## Discussion

In this study, we purified mitochondria from porcine hearts and utilized cryo-ET and related approaches to elucidate the *in-situ* structure of mammalian PDC. Our results show the intrinsic flexibility of the PDC on both diameter and peripheral organizations. In mitochondrial PDC, E1 scatters around the icosahedral PDC core, while E3 only locates along the 5-fold axis (Fig. 2f-g). This could be attributed to that the spacious pentagonal channel, compared to that near the 2-fold and 3-fold axes, could facilitate the mobility of the E3 outer motif more effectively, enabling its active involvement in the oxidation process of dihydrolipoyl moieties associated with the five adjacent E2 trimers. Two normal distributions of the number of E1 and E3 in each mitochondrial PDC were observed (Fig. 2c), suggesting that there may be distinct subpopulations or heterogeneity within the PDC population and an additional regulation to promptly respond to different cellular states or nutrition conditions.

The exact number of E2 and E3BP constituting the 60-mer core of mammalian PDC is still under debate. A "substitution model" for mammalian E2:E3BP favors 40:20^54^ or 48:12^23^. Due to the resolution limit, we are unable to discriminate the densities of E3BP and E2 within our *in-situ* PDC core structure. However, since E3 specifically binds on the E3BP-linked PSBD, our results reveal up to 12 E3 dimers in the mitochondrial PDC, implying that the mammalian PDC core scaffold is likely composed of 48 E2 and 12 E3BP. In addition, although one PDC core could bind up to 48 E1 and 12 E3 multimers, the actual numbers of E1 and E3 multimers are typically far from saturation (Fig. 2c), similar as *M. Smolle et.al* proposed^43^. Such structural features might also be true for prokaryotic PDCs, which could function properly without fully saturating all the binding sites of E1 and E3^22^, potentially allowing various substitutions to emerge during evolution. The dynamic movement of the N-terminally-located, E2-linked LDs plays a pivotal role in the catalytic process of PDC. These LDs, functioning as swinging arms, facilitate the transferring of their prosthetic groups to the active sites of all three enzymes throughout the catalytic cycle. Here, we captured the dynamic interactions of LDs with both E1 (Fig. 3) and E2 (Fig. 4). These interactions primarily rely on electrostatic interactions (Fig. 3i-j and Fig. 4j-k). We thus propose that the acidic patches on the LD serve as a general binding site for its interactions with different components of PDC. The reported LD position relative to E2 in the resting state of PDC in *Escherichia coli* is different from our observations (Extended Data Fig. 6f), whereas the position of the lysine residue (K245 in *Escherichia coli* LD and K259 in porcine LD) barely changes (Extended Data Fig. 6g). This confirms that the E2-LD interactions are highly dynamic and suggests that LD might be able to deliver the substrate-carrying prosthetic group into the E2 active site from various directions.

Previous study^18^ and our present work show that the binding sites of E1 and E3 are not necessarily fully saturated for a PDC in mitochondria to be functional. The E2 core associated with one peripheral E1 heterotetramer and one E3 dimer is sufficient to accomplish the catalysis^22^. Consequently, the more peripheral enzymes, the more LD substrate binding sites, and the higher the catalytic efficiency. Our results suggest that while the number and distribution of the peripheral E1 and E3 multimers vary substantially, the mitochondria may still normally carry out energy metabolism. The variation of E1 or E3 quantity may also function to regulate the catalytic efficiency of PDC in addition to other ways of regulation, and may be controlled by physiological conditions and metabolic demands of mitochondria.

We therefore proposed an additional regulation model for the catalytic performance of mammalian PDC. In low-energy demand state, several E1 and E3 enzymes are adequate for the catalytic reactions (Fig. 5a). Although each E2 possesses two LDs, only one LD is required for substrate transfer since there is a limited number of E1/E3 active sites (Fig. 5b-d). E1 captures and decarboxylates a pyruvate and transfers it to a mobile LD (Fig. 5b). Then, the substrate is transferred by the LD to an active site on E2 to produce acetyl-CoA (Fig. 5c). Subsequently, the lipoyl moiety is transferred to a catalytic site of E3 by the LD and reoxidized to complete one reaction (Fig. 5d).

**Fig. 5.**
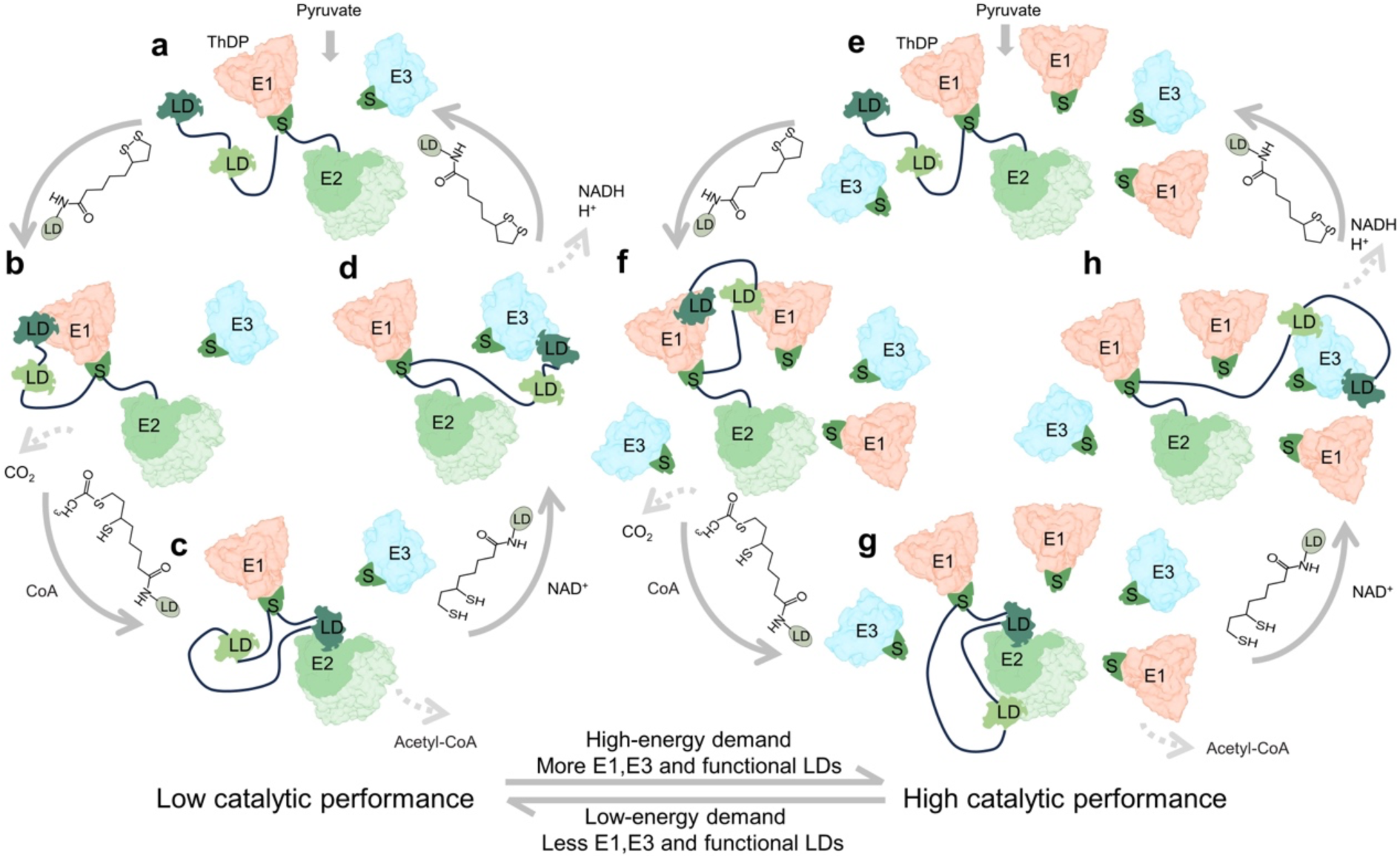
A catalytic performance regulation model of the PDC. **a-d,** Schematic of PDC reaction with low catalytic performance. In low energy demand, limited amount of E1 and E3 bind to the PDC core. Each E2 has two LDs, but only one of the LDs functions as swing arm to carry out substrate transfer among E1, E2 and E3. **e-h,** Schematic of PDC reaction with high catalytic performance. With increasing energy demand, more E1 and E3 bind to the PDC core and provide more activating sites. Both LDs of the E2 are required to function as swing arms for substrate transfer to increase the catalytic performance. All E1 and E3 bind on the PSBD connected with the E2 core by flexible linkers. Some linkers are omitted for simplicity. LD: lipoyl domain, S: PSBD.

In a high-energy demand state, more E1 and E3 multimers would be assembled on the PDC core to provide more active sites, and both LDs of the E2 subunit are required to function simultaneously to achieve higher efficient transfer of substrate (Fig. 5e-h). Both LDs interact with E1 subunits simultaneously (Fig. 5f) and transfer acetyl groups, facilitating reactions at two catalytic sites on the E2 heterotetramers (Fig. 5g), then transfer the resulting reduced lipoyl moiety to two catalytic sites on E3 (Fig. 5h), thereby completing multiple reactions simultaneously to improve the catalytic efficiency. This high-performance catalytic process continues until the energy demand of the mitochondrion decreases. Then part of the E1 and E3 dissociate from the E2 core and the PDC reverts to the state where only one LD is involved in the reaction. Previous studies have postulated two different mechanisms for the higher-order interaction of the components of OADH complexes: the ‘‘multiple random coupling’’ (MRC) mechanism^55^ and the ‘‘division-of-labor’’ (DOL) mechanism^45,56^. The model we proposed favors the MRC mechanism in which the overall activity of the PDC is influenced by redundancies and random processes. Considering the similarity among members of the OADH family, the regulation mechanism proposed here is likely applicable to all members of the OADH family.

The regulation of the mammalian PDC is crucial for maintaining glucose homeostasis during feeding and fasting states. It has been reported that the catalytic reaction of PDC is regulated through the end-product inhibition by acetyl-CoA and NADH^57,58^, and through the reversible phosphorylation and dephosphorylation of the E1 subunit by PDK and PDP^3,59,60^. Acetyl-CoA/CoA and NADH/NAD+ ratios act as effectors in feedback inhibition by influencing the redox and acetylation status of the lipoamide cofactors. While PDK and PDP inhibit PDC activity by phosphorylation of the serine residues on E1. Since the (de-)phosphorylation regulation directly relates with the enzyme activity of E1, we suspect such (de-)phosphorylation of the E1 subunit might affect the (dis-)assembly of E1 and E3 to regulate the catalytic performance of PDC. It may be helpful to select periods of high and low expression of PDP and PDK to obtain samples and carry out structural analysis to verify this hypothesis.

In conclusion, we obtained the *in-situ* structure of mammalian PDC which implies the dynamic substrate transporting by the LDs between different enzymes. The location and quantity of peripheral proteins as well as the interface at which they interact with the PDC core are structurally described. Overall, our results provide profound insights into the structure and regulation of OADH family complexes.

## Methods

### Isolation of mitochondria from porcine heart

All the following procedures were carried out on ice or at 4°C. Fresh porcine heart was obtained from the slaughter house and transported to the laboratory as soon as possible. The heart muscles were washed twice with Milli-Q water. 720g of heart muscles were isolated and cut into small cubes and suspended in 1000 ml of buffer A (0.25Msucrose, 10mM potassium phosphate, 0.1mM EDTA, pH 7.6). These tissues were homogenized in a large-capacity blender for 5 min. The homogenate was centrifuged in l-liter bottles at 2000 g for 10 min. The supernatant fluid was strained through eight layers of cheesecloth. The mitochondrial fraction was collected by centrifugation at 22,000 g for 17 min. The Crude mitochondria was washed twice with 200 ml 0.02 M potassium phosphate, pH 7.0 and paste was collected by centrifugation at 22,000 g for 10 min. The crude extracted mitochondria were prepared for further purification of PDC or preparation of cryo-ET samples.

### PDC purification

The crude mitochondria were suspended in 60 ml 0.02 M potassium phosphate (pH 7.0). 5 M NaCl and Cooktail were added into the suspension at a ratio of 1:100, and the pH was adjusted to 6.4. The suspension was disrupted twice at 276 bar with a cell disruptor (JNBIO) equipped with a cooling system. The mitochondria lysate was centrifuged at 30,000 g for 30 min, and the supernatant was added 10 mM MgCl_2_ and PEG precipitated in 5% (w/v) PEG 6,000 for 10 min. The precipitate was collected by centrifugation at 30,000g for 20 and resuspended in 25 ml buffer B (50 mM MOPS-KOH buffer, pH 7.0, 0.02 mM thiamin pyrophosphate, 1 mM MgCl_2_, 2mM DTT). The suspension was stored 4h in a refrigerator and then centrifuged at 40,000 g for 20 min before discarding the precipitates. To the clear supernatant fluid was added 2 mM EGTA. After 20 min the solution was loaded onto 10ml 35% (w/v) sucrose cushions centrifuged at 105,000 g for 3.5 h in a Beckman type SW32 rotor. The precipitate is resuspended with buffer B supplemented with 2 mM EGTA. After 20 minutes, the solution was added to a 10% to 40% sucrose gradient configured by buffer C (50 mM potassium phosphate buffer, pH 7.0, 0.02 mM thiamin pyrophosphate,1 mM MgCl_2_, 1 mM NAD^+^, 0.1 mM EDTA, 0.4 mM DTT)) and centrifuged at 144,000 g for 2.5 h using a SW41Ti rotor with a Beckman Coulter Optima XPN-100 centrifuge. The gradients were fractionated and assessed by SDS-PAGE and negative staining, PDC around the sucrose density of 15%. The samples were concentrated to 0.4 mg ml^-1^ by using a membrane concentrator with a 100 kDa cut-off for biochemical and cryo-ET analyses.

### Biochemical analysis of PDC

Prepare a 12% polyacrylamide gel and assemble the electrophoresis tank. Load 20 μl of PDC protein sample and 5 μl of protein marker onto the gel for polyacrylamide gel electrophoresis (SDS-PAGE).

Purified PDC obtained by purification method was concentrated to 30 μl using a 100 kDa concentrator. Protein precipitation was performed using TCA precipitation method. The precipitated protein was resolubilized by adding a resolubilization solution (8 M urea,100 mM Tris-HCl, pH 8.5), and the protein concentration was determined using the BCA method. Equal amounts of protein were taken and adjusted to the same volume. TCEP (tris(2-carboxyethyl)phosphine) and CAA (chloroacetamide) were added for reduction and alkylation reaction at 37 ℃ for 1 hour. The reduced and alkylated samples were diluted with 100 mM Tris-HCl solution to reduce the Urea concentration to below 2 M. Trypsin was added at a 1:50 ratio of enzyme to protein and incubated overnight at 37 ℃ with shaking for enzymatic digestion. The next day, the digestion reaction was terminated by adding TFA, and the supernatant was subjected to desalting using an SDB-RPS desalting column. After vacuum drying, the samples were stored at -20 ℃. Mass spectrometry analysis was performed using an ion mobility-quadrupole-time-of-flight mass spectrometer (timsTOF Pro) from Bruker. Sample injection and separation were performed using an UltiMate 3000 RSLCnano liquid chromatography system coupled online with the mass spectrometer. Peptide samples were injected and trapped on a Trap column (75 μm × 20 mm, 2 μm particle size, 100 Å pore size, Thermo), followed by elution onto an analytical column (75 μm × 250 mm, 1.6 μm particle size, 100 Å pore size, ionopticks) for separation. The analysis gradient was established using two mobile phases (mobile phase A: 0.1% formic acid in H2O and mobile phase B: 0.1% formic acid in ACN). The flow rate of the liquid phase was set to 300 nL/min. Peptides were introduced into the mass spectrometer via CaptiveSpray nanospray ion source for DDA scanning. TIMS function was enabled, and PASEF scanning mode was used. Each scan cycle consisted of 1.1s, including one MS1 scan and 10 PASEF MS/MS scans, with each PASEF MS/MS scan generating 12 MS/MS spectra. The mass spectrometry data were analyzed using MaxQuant (V2.0.1) software with the Andromeda database search algorithm. The protein reference database used for the search was the Sus_scrofa protein reference database from Uniprot (2023-01-09, containing 46,139 protein sequences). The main search parameters were as follows: variable modifications - Oxidation (M), Acetyl (Protein N-term); fixed modification - Carbamidomethyl (C); enzyme - Trypsin/P; primary mass tolerance - 20 ppm in the first search and 4.5 ppm in the main search; secondary mass tolerance - 20 ppm. The search results were filtered at a 1% FDR (false discovery rate) threshold at the protein and peptide levels. Protein entries corresponding to reverse sequences, contaminating proteins, and peptides with a single modification were removed, and the remaining identification information was used for further analysis.

### Preparation of cryo-ET samples

An aliquot of a 3μl purified PDC sample was applied to a holey carbon grid covered with graphene-oxide (Quantifoil R1.2/1.3, Au, 300 mesh). After waiting for 60 s, the grids were blotted for 3.5 s at a humidity of 100% and 4°C and plunge-frozen in liquid ethane using a Vitrobot (Thermo Fisher Scientific).

The crude extracted mitochondria was rinsed gently with 1 ml of 0.02 M potassium phosphate. An aliquot of a 5 μl sample was applied to a glow-discharged holey carbon grid (Quantifoil R2/2, Au, 400 mesh). After waiting for 60 s, the grids were blotted for 7 s at a humidity of 100% and 22 °C and plunge-frozen in liquid ethane using a Vitrobot.

### Cryo-ET data collection and tomogram reconstruction

The frozen-hydrated samples were transferred to a 300 kV Titan Krios electron microscope (Thermo Fisher Scientific) equipped with a GIF energy filter and a Falcon 4 direct electron detector (Thermo Fisher Scientific). All images were recorded at ×81,000 magnification with a pixel size of 1.5 Å at the specimen level. PaceTomo script^61^ was used with the SerialEM software to collect tilt series at -2 to -4 µm defocus (dose-symmetric collection theme, start from 0°, group size 2), with accumulative dose of ∼90–100 e−/Å^2^ distributed over 35 images and covering angles from −51° to 51°, with a tilt step of 3°.

All recorded images were first motion-corrected by Relion-4.0^62^ and then stacked by IMOD^63^ and aligned by AreTomo^64^. Gctf^65^ was used to determine the defocus of each tilt image. For subtomogram averaging using i3, the ‘ctfphaseflip’ function in IMOD was used to do the contrast transfer function (CTF) correction for the tilt images. Tomograms were reconstructed by weighted back-projection (WBP) or simultaneous iterative reconstruction (SIRT) methods using IMOD or tomo3D with the CTF-corrected aligned stacks.

### Subtomogram averaging and corresponding analysis

All PDC core particles were first manually picked from 6x binned SIRT reconstructed tomograms. Then subtomograms of PDC particles were cut from the 4x binned WBP reconstructed tomograms and aligned by i3^66^.

Refinement of the PDC core structure: the subtomogram averaging pipeline in Relion-4.0 was used for high-resolution structure determination. The coordinates and Euler angles of each PDC particle generated from the previous i3 alignment step were first transferred to Relion4.0 by a homemade script package. 3D alignment and classification were generated under binning 4x and 2x, followed by local refinement and 2 times of CTF and frame refinement for unbinned data, resulting in the 4.3 Å resolution PDC core structure.

Local refinement and classification of E2 subunit: each PDC core consists of 60 E2 subunits. After the alignment of PDC core, the position of each E2 subunit was calculated using the “symmetry expansion” and “shift center” functions in Relion-4.0 (60 E2 subunits corresponding to each PDC core). Then Relion-3.1 was used to do 3D alignment and classifications for all E2 subunits, under binning 4x and 2x, based on the previously described protocol^67^ with some minor modifications. Duplicates were removed for particles with distance less than 3 nm between their centers. A spherical mask was applied adjacent to the E2 subunit to generate the classification results shown in Fig. 3b-d. Afterwards, the multi-reference alignment (MRA) and classification in i3 were applied to further confirm the classification result.

Refinement and classification of E1 and E3 multimers: 1) we speculate the densities surrounding the PDC cores in raw tomograms (Fig. 1c and Extended Data Fig. 1a) are E1 and E3 multimers. Therefore, we manually picked ∼10k particles of these densities from the 6x binned SIRT reconstructed tomograms, then the corresponding subtomograms were cut from the 4x binned WBP reconstructed tomograms, followed by 3D alignment and classification using i3 to get an initial model. In the initial model, part of the icosahedral PDC core is still visible, with strong densities roughly locating along its C2, C3 and C5 axis and the distance between the densities and PDC core is ∼20 nm. 2) Using the PDC core orientations and positions from Relion-4.0, we run “symmetry expansion” (icosahedral symmetry) for each PDC core, then shifted the positions to ∼20 nm away from the core center. We thus got 60 positions corresponding to each PDC core. Those positions were used as particle centers to cut subvolumes from the WBP reconstructed tomograms. The volume size was set to ∼40 nm to make sure there are enough overlaps between different subtomograms, so that we did not miss any possible E1 or E3 particles. 3) The subtomograms were aligned to the initial model generated from i3, followed by 3D classifications. E1 and E3 multimers are identified by comparing the density shape with the reported E1 and E3 structures. See also Extended Data Fig.1 for subtomogram averaging workflow.

### Mitochondrial volume size calculation

Using the IMOD software^63^, we manually draw the area of each mitochondrion in all tomograms and obtained the area size based on the IMOD contour information. We then rotate the tomogram along the Z-axis, measure the height of the mitochondria, and calculate the volume size of each mitochondrion by multiplying the area size with the height.

### Model Fitting and Visualization

Atomic models (PDB accession code E2: 7uom, E1 binding PSBD: 6cfo, 1w88, E3: binding PSBD: 1zy8, PSBD: 1zwv) were rigidly fitted to the corresponding densities using the Fit in Map tool^68^ in UCSF chimera or ChimeraX. EMAN2 was used for segmentation of the mitochondrial membranes and cristae. ArtiaX^69^ was used for mapping the PDC complexes and ATP back to the raw tomograms. UCSF Chimera^68^ and ChimeraX^70^ were used for rendering the graphics.

## Data and materials availability

Cryo-ET subtomogram averages were deposited in Electron Microscopy Data Bank under accession codes: E2/E3BP core (Fig. 4b): EMD-38712, E1-PSBD (Fig. 3b): EMD-38711, E1-PSBD-LD1 (Fig. 3c): EMD-38709, E1-PSBD-LD2 (Fig. 3d): EMD-38710, E2-LD1 (Fig. 4c): EMD-38713, E2-LD2 (Fig. 4d): EMD-38714, E2-Linker (Fig. 4e): EMD-38715, E3-PSBD (Fig. 3k): EMD-38716. All data needed to evaluate the conclusions in this paper are present in the paper and the supplementary information.

## Acknowledgements

We thank Lingyun Wu, Chenyu Yang and Menghan Zhang in the Center of Cryo-Electron Microscopy (CCEM), Zhejiang University for their technical assistance on 200kv and 120kv cryo-electron microscopy. We thank Deyue Xu from the Laboratory Animal Center, Zhejiang University for providing fresh porcine heart in pre-experiment. We thank Shuaiqi (Phil) Guo and Donghyun (Raphael) Park for critical reading of the manuscript. This work was supported by the National Key Research and Development Program of China (2018YFA0507700 to X.Z.) and National Natural Science Foundation of China (32200980 to Y.C.).

## Author contributions

X.Z. initiated the project. X.Z. and Y.C. supervised the research. C.W., C.M., H.W., Y.W., S.A. and Q.H. performed the sample preparation and characterization. Y.C., C.W. and S.C. collected the cryo-ET data. Y.C. and C.W. processed the cryo-ET data and reconstructed the cryo-ET density map. C.W. and Y.C. prepared figures. M.C., C.W., S.C. and J.C. performed the AlphaFold2 prediction and sequence alignment. C.Y., J.X, Y.J. and Z.J. assisted in the sample preparation and data analysis. Y.C., X.Z., C.W., C.M. and H.G. wrote the manuscript. All authors discussed and commented on the results and the manuscript.

## Additional Information

**Supplementary Information** is available for this paper.

**Correspondence and requests for materials** should be addressed to Yunjie Chang or Xing Zhang.

## Extended Data

**Extended Data Fig. 1.**
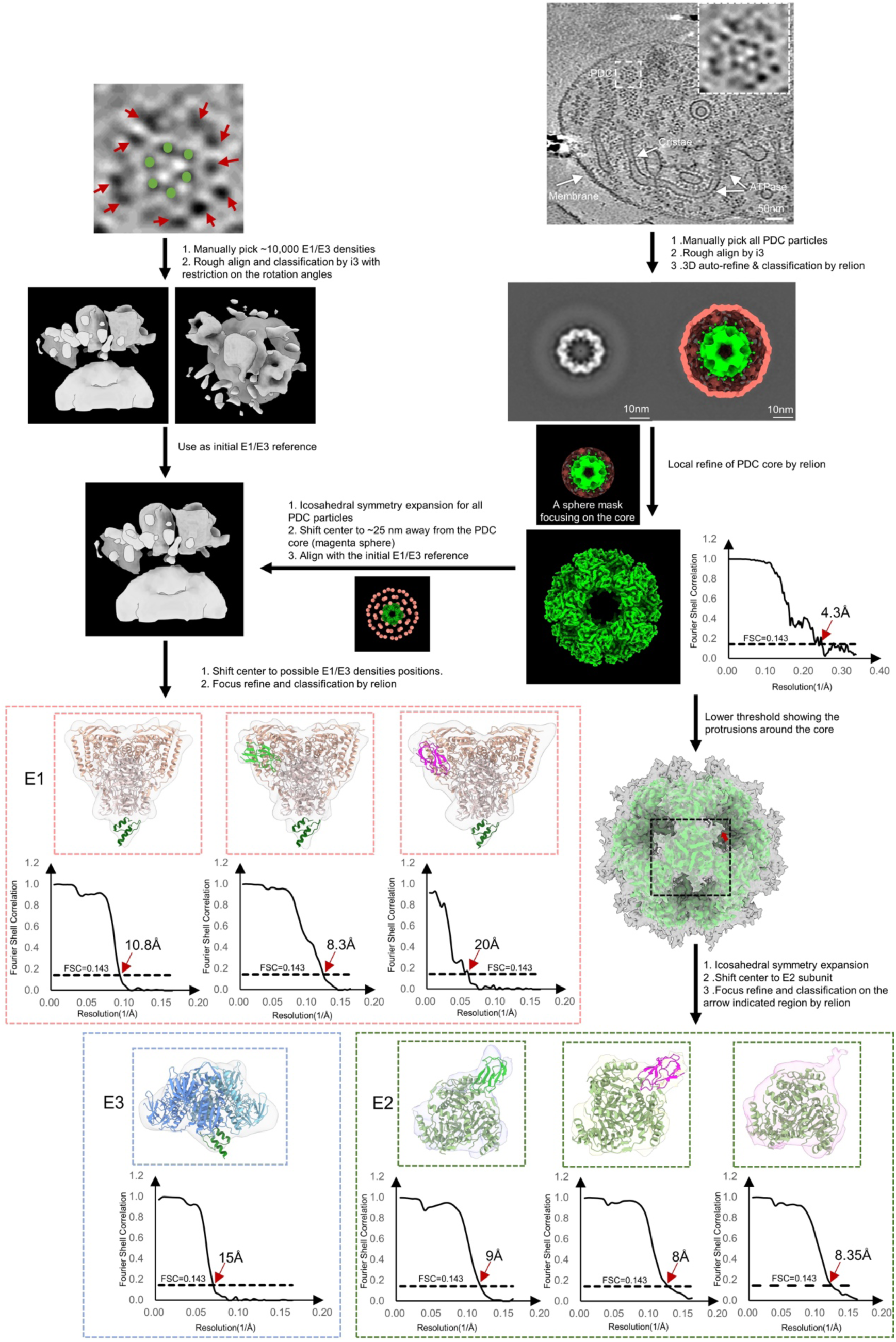
Subtomogram averaging workflow. Flow chart of the subtomogram averaging processing. The PDC particles were manually picked from the tomograms and then averaged by i3 or Relion. The PDC core and peripheral densities are then local refined and classified to improve the resolution and analyze the interactions between different components. Refer to Materials and Methods for details of the processing.

**Extended Data Fig. 2.**
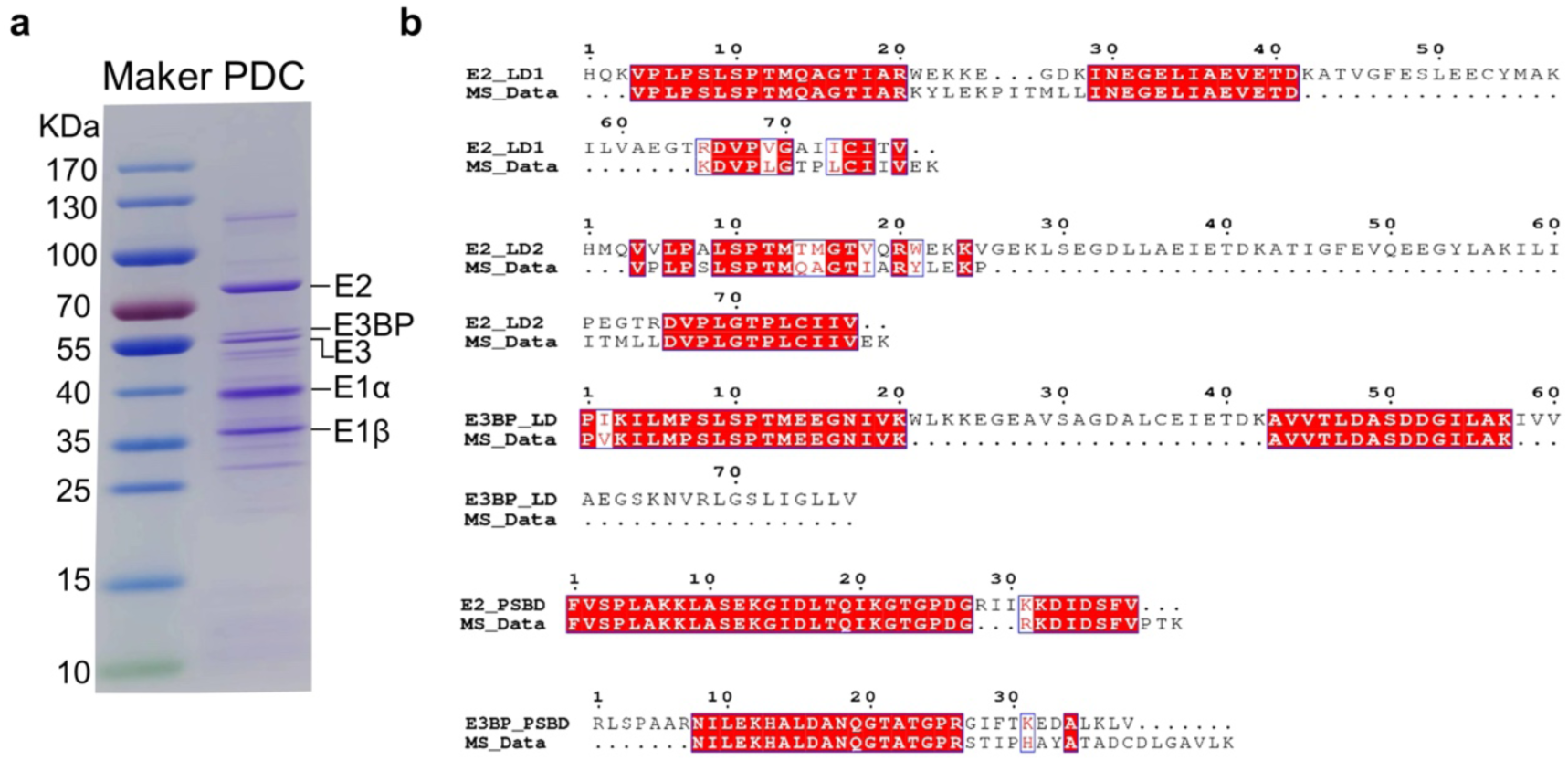
SDS-PAGE analysis of the purified PDC and mass spectrometry analysis of the LD and PSBD. **a,** The SDS-PAGE analysis detected the purified PDC. The resulting bands include the major components of the PDC: E2, E3BP, E3, E1α, E1β, lane1: pre-stained marker; lane 2: purified PDC sample. This data is repeated more than three times, and all resulted in the same results. **b,** The mass spectrometry data analysis comparison of peptides contained in DLAT (E2) and PDHX (E3BP) with LD and PSBD and same amino acids are represented in red.

**Extended Data Fig. 3.**
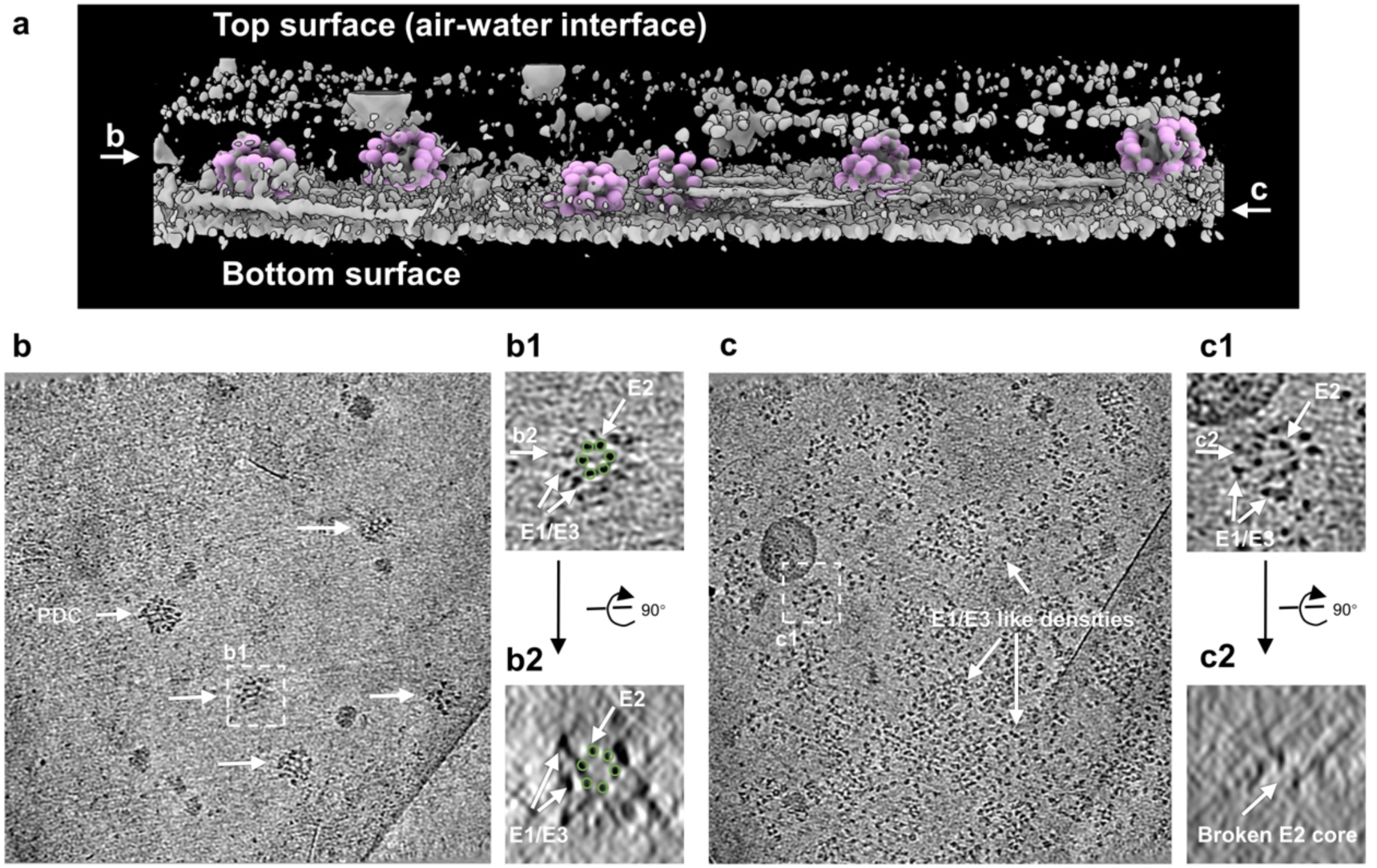
Cryo-ET investigations of the purified PDC. **a,** 3D surface view of a representative cryo-electron tomogram of the purified PDC sample (viewing roughly along the y axis). The densities of six unbroken PDC particles are shown in pink spheres. **b,** A section (viewing along z axis) close to the center of the tomogram shown in **a**. Five PDC particles are indicated by arrows. The enlarged view of a PDC is shown in **b1**. **b2** shows the side view of the same PDC. The E2 core is clearly visible in both **b1** and **b2** (indicated by green circles). The densities surrounding the E2 core are supposed to be E1 or E3. **c,** A section (viewing along the z axis) close to the bottom surface of the tomogram shown in **a**. Multiple PDCs like densities are observed in this tomogram section and one example is shown in **c1** (top view) and **c2** (side view). The E1, E2 and E3 densities are clearly visible in top view **c1**. In the side view **c2**, however, the E2 core appears to be broken and the E1 and E3 are almost invisible. Instead, E1 and E3 like densities scattering around the bottom surface of the tomogram.

**Extended Data Fig. 4.**
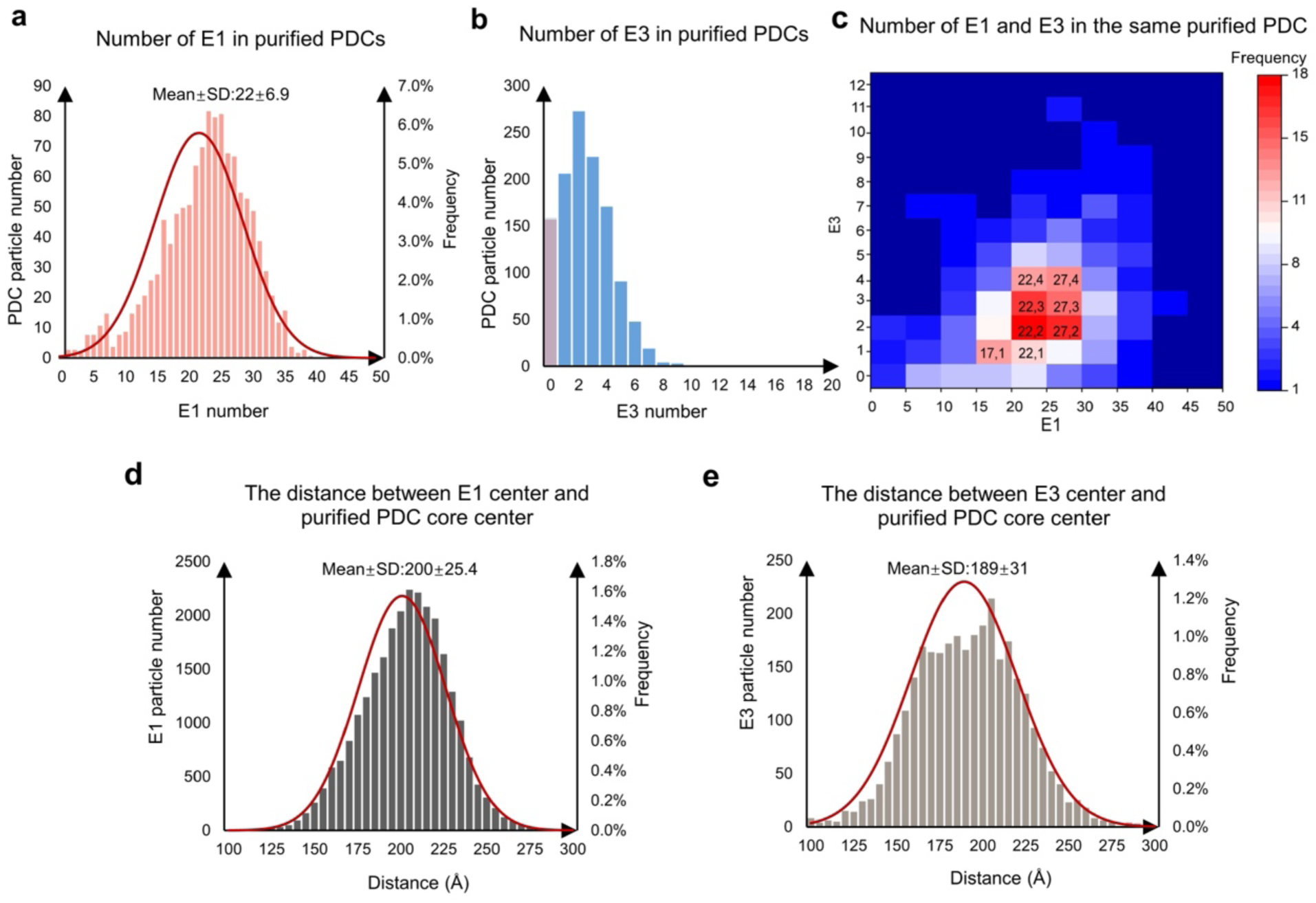
The numbers and spatial arrangements of E1 and E3 multimers in purified PDC. **a,** The number of E1 heterotetramer in purified PDCs. b, The number of E3 dimer in purified PDCs. c, The number of E1 and E3 multimers in the same purified PDC. d, Statistics on the distance between the E1 heterotetramer center and the PDC core center. e, Statistics on the distance between the E3 dimer center and the PDC core center.

**Extended Data Fig. 5.**
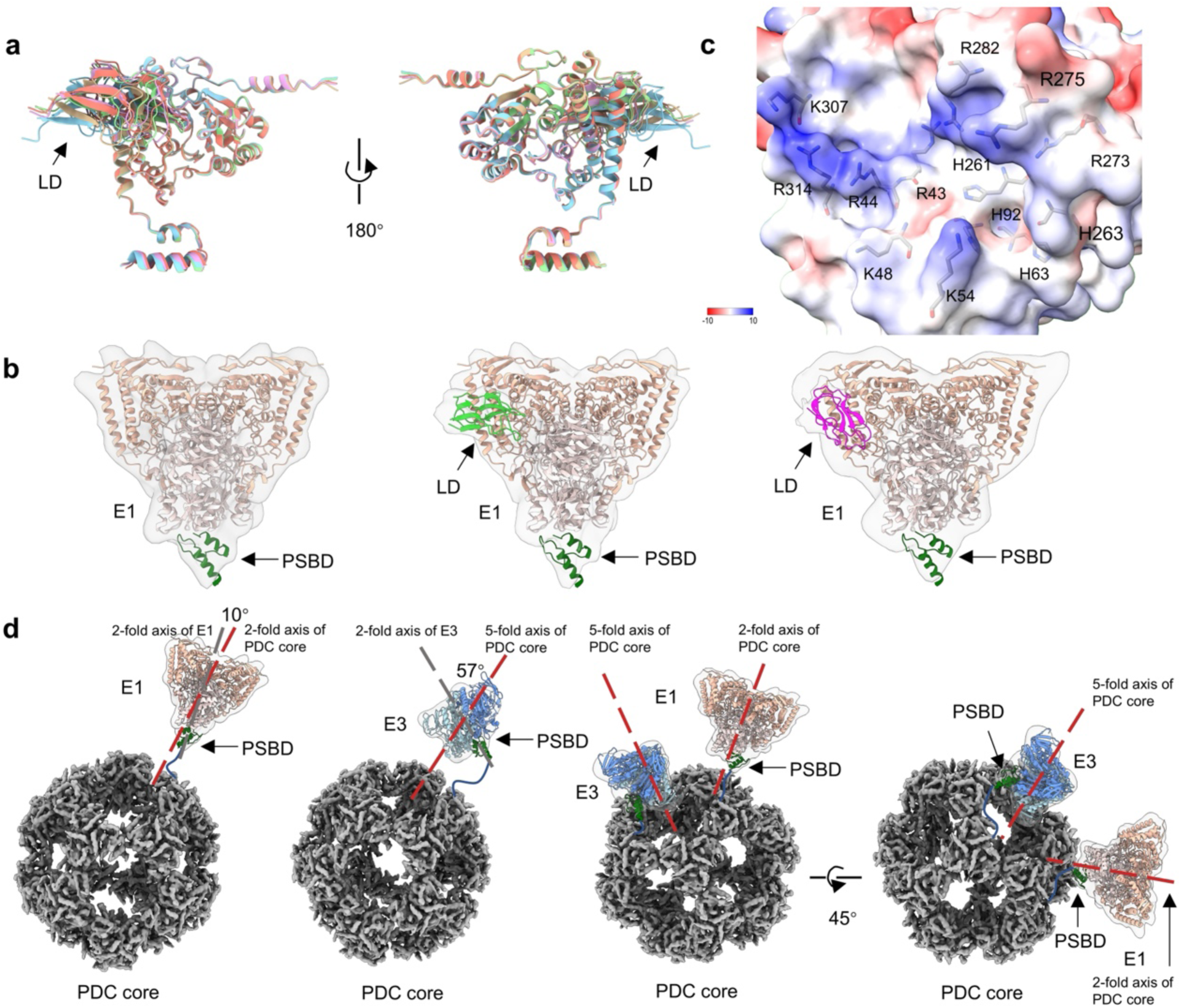
The interaction and orientation between E1 and E3 with the LD and PSBD of the PDC core. **a,** AlphaFold2 predicts five distinct conformations for the binding of the C chain of E1 with LD. LD locates near the E1 N-terminal region. **b,** Class averages of E1 heterotetramer, with PSBD fitting in opposite orientations compared to those shown in Fig. 3b-d. **c,** Analysis of the electrostatic potential of E1 at the LD-E1 interaction interface, showing the location of basic amino acids. d, The positions and orientations of E1 and E3 multimers relative to the PDC core are different. E1 heterotetramer binds with PSBD in a head-on manner towards the core, while the E3 dimer binds with PSBD at a certain angle relative to the core.

**Extended Data Fig. 6.**
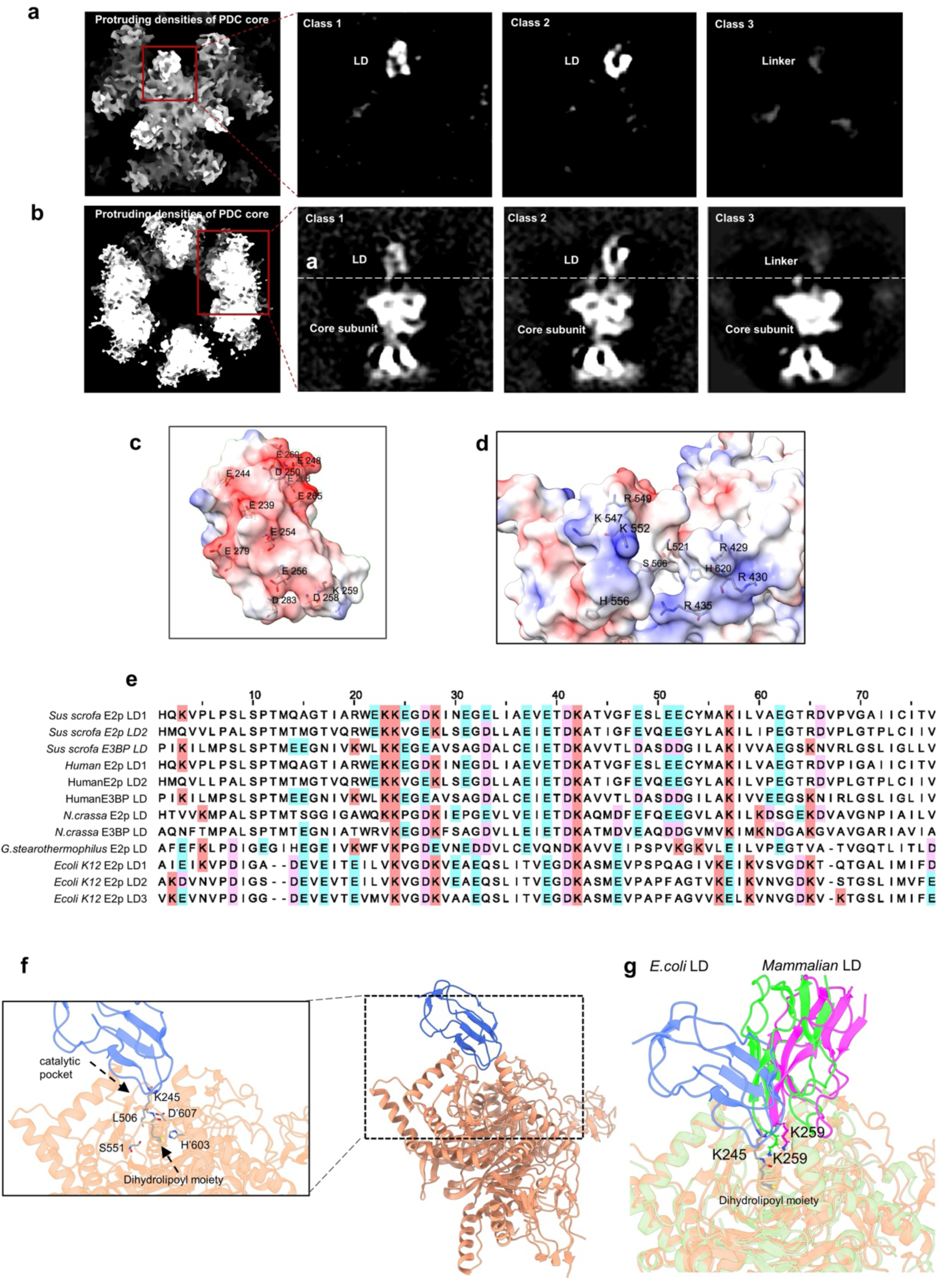
Subtomogram average structures of E2 and the charged potential energy analysis at the interfaces between E2 and LD. **a,** The top view of 3D classification of protruding densities of PDC core, divided into three classes. Class 1 and Class 2 are referred to as LD (Low Density), while Class 3 is referred to as linker. **b,** The side view of 3D classification of protruding densities of PDC core. **c,** Acidic patches on the surface of LD, showing the location of acidic amino acids. **d,** Electrostatic potential of E2 at the LD-E2 interaction interface, showing the location of basic amino acids. **e,** The LD sequences of different species were aligned, with acidic amino acid E highlighted in blue, D highlighted in pink, and conserved amino acid K highlighted in light red. **f,** The E2-LD interaction interface in resting state PDC of *Escherichia coli* (PDB 7b9k). **g,** Comparison between the E2-LD interaction sites of mammalian PDC from current work and the resting state *Escherichia coli* PDC. The catalytic pocket of E2 and LD of *Escherichia coli* PDC (Fig. 2c and d) are shown in orange and blue, respectively. The catalytic pocket of E2 of mammalian PDC is shown in dark sea green, and the mammalian LDs with two different conformations are shown in lime and magenta, respectively.

**Extended Data Table 1.**
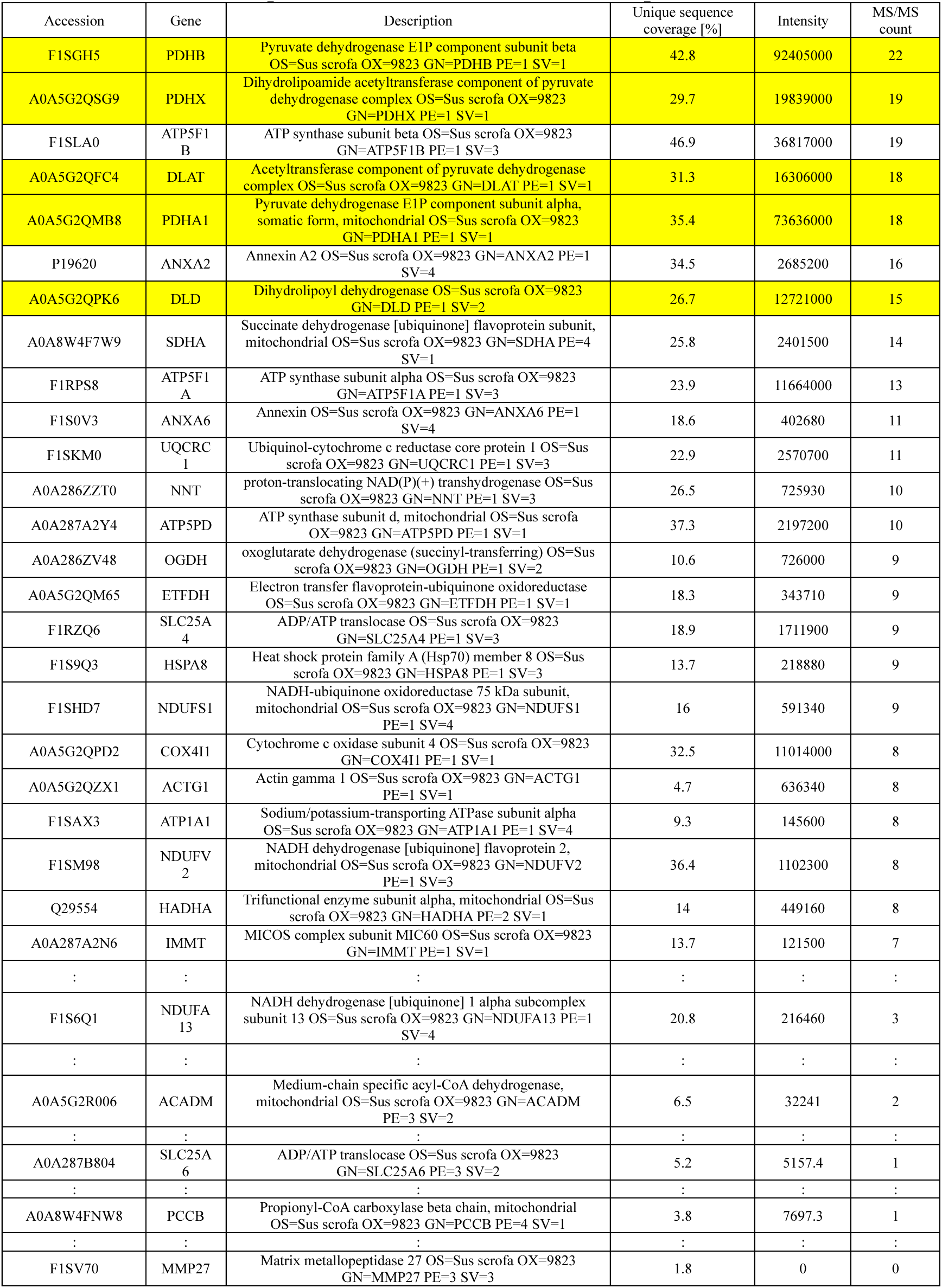
Mass spectrometry analysis. Mass spectrometry analysis was performed on the purified protein solution of PDC, with the main components of PDC highlighted in yellow. The complete mass spectrometry analysis data can be found in the source data file. The Excel file contains multiple worksheets and includes descriptions for each column header.

**Extended Data Table 2.** Number of tomograms and particles used for subtomogram averaging analysis.

|  | Tomograms | PDC particles | All E1 | E1-PSBD (Fig. 3b) | E1-PSBD-LD1 (Fig. 3c) | E1-PSBD-LD2 (Fig. 3d) | E2-LD1 (Fig. 4c) | E2-LD2 (Fig. 4d) | E2-Linker (Fig. 4e) | E3 (Fig. 3k) |
| --- | --- | --- | --- | --- | --- | --- | --- | --- | --- | --- |
| **Purified PDC** | 256 | 1,209 | 27,156 | 3,862 | 22,951 | 343 | 16572 | 29289 | 26679 | 3,120 |
| **Mitochondrial PDC** | 122 | 805 | 17,038 | 3,097 | 13,755 | 186 | 10552 | 20811 | 16929 | 2,977 |
| **Total** | 378 | 2,014 | 44,194 | 6,959 | 36,706 | 529 | 27124 | 50100 | 43608 | 6,097 |

**Extended Data Table 3.**
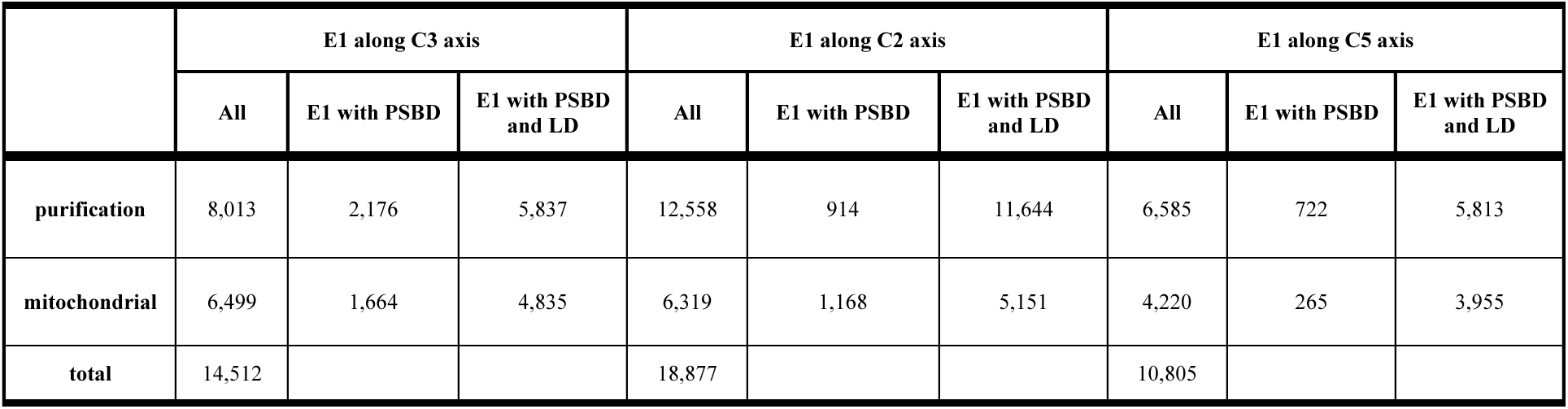
The quantity distribution of E1 heterotetramer above the 2-, 3-, and 5-fold axes of the PDC core.

|  | E1 along C3 axis |  |  | E1 along C2 axis |  |  | E1 along C5 axis |  |  |
| --- | --- | --- | --- | --- | --- | --- | --- | --- | --- |
|  | All | E1 with PSBD | E1 with PSBD and LD | All | E1 with PSBD | E1 with PSBD and LD | All | E1 with PSBD | E1 with PSBD and LD |
| **purification** | 8,013 | 2,176 | 5,837 | 12,558 | 914 | 11,644 | 6,585 | 722 | 5,813 |
| **mitochondrial** | 6,499 | 1,664 | 4,835 | 6,319 | 1,168 | 5,151 | 4,220 | 265 | 3,955 |
| **total** | 14,512 |  |  | 18,877 |  |  | 10,805 |  |  |

